# Applying Mondrian Cross-Conformal Prediction to Estimate Prediction Confidence on Large Imbalanced Bioactivity Datasets

**DOI:** 10.1101/116764

**Authors:** Jiangming Sun, Lars Carlsson, Ernst Ahlberg, Ulf Norinder, Ola Engkvist, Hongming Chen

**Author notes:** Corresponding Author Hongming Chen.

## Abstract

Conformal prediction has been proposed as a more rigorous way to define prediction confidence compared to other application domain concepts that have earlier been used for QSAR modelling. One main advantage of such a method is that it provides a prediction region potentially with multiple predicted labels, which contrasts to the single valued (regression) or single label (classification) output predictions by standard QSAR modelling algorithms. Standard conformal prediction might not be suitable for imbalanced datasets. Therefore, Mondrian cross-conformal prediction (MCCP) which combines the Mondrian inductive conformal prediction with cross-fold calibration sets has been introduced. In this study, the MCCP method was applied to 18 publicly available datasets that have various imbalance levels varying from 1:10 to 1:1000 (ratio of active/inactive compounds). Our results show that MCCP in general performed well on cheminformatics datasets with various imbalance levels. More importantly, the method not only provides confidence of prediction and prediction regions compared to standard machine learning methods, but also produces valid predictions for the minority class. In addition, a compound similarity based nonconformity measure was investigated. Our results demonstrate that although it gives valid predictions, its efficiency is much worse than nonconformity measures obtained from supervised learning.

## INTRODUCTION

To address the increasing drug development costs and reduced productivity faced by the pharmaceutical industry, QSAR/QSPR (Quantitative Structure-Activity/Property Relationship) models have gained popularity for predictions of biological activities and physicochemical properties as well as for in silico screening of large number of compounds. Informed decisions based on predictions from a QSAR model are frequently confounded by a poor understanding of the confidence of the prediction for the compound of interest. These computational models are not guaranteed to give equally accurate predictions in all of the chemical space of interest. In other words, the QSAR models have limited applicability domain (AD). The AD refers to the chemical space where the property can be predicted by the model with high confidence. An assumption of the AD concept is that the further away a molecule is from a QSAR model’s AD (according to a given measure), the less reliable the prediction is.

A number of metrics^1-2^ have been proposed in the literature to define the AD, e.g. “distance/similarity to model”, “bagged variance” and “reliability indices”. The most common type is distance-to-model metrics^3-5^ that measure the distance between a test compound and the training set for the model to estimate the “closeness” between the test compound and the training set. This is done by calculating the distance between the used descriptors according to a specified metric. Alternative approaches include defining regions of the descriptor space with different levels of reliability^6-7^ and, subsequently, assessing the prediction error using sensitivity analysis that samples or perturbs the composition of the training set to estimate a distribution of predictions^8^. Recently, another type of AD measurement was proposed by building an additional error model to assess the prediction reliability^9-10^. Benchmark studies^2^, ^11-12^ on various AD metrics have previously been performed. Toplak et al^12^ showed that methods of reliability indices were sensitive to dataset characteristics and to the regression method used in building the QSAR model. Most of these AD metrics lack a rigorous scientific derivation. Their correlation with prediction confidence is only empirically validated and might therefore be dataset dependent. In practice, what an experimentalist would like to know is if a prediction falls in a given prediction interval with a certain confidence, for instance a prediction with 80% or 95 % confidence.

Conformal prediction^13-15^ is a method for using known data to estimate prediction confidence for new examples. It has recently been proposed to address insufficiencies of earlier AD metrics in the QSAR domain^16-17^. Our previous studies^18-19^ have shown that conformal prediction provides a rational and intuitive way of interpreting AD metrics as prediction confidence with a given confidence level. Most AD estimates can actually be seamlessly used within the conformal prediction framework. Conformal prediction is also a rigorously defined concept within statistical learning theory. To deal with imbalanced data sets, the Mondrian conformal prediction (MCP) was introduced^20^. It divides data according to their label where a separate significance level is set for each class. Therefore, MCP can guarantee validity for each class. MCP have been applied to diagnose bovine tuberculosis^21^.

In the current study, a novel conformal prediction protocol, Mondrian cross-conformal prediction (MCCP) has been used to estimate the confidence of predictions. It has been applied to several large cheminformatics datasets with various levels of imbalance between number of active and inactive compounds. The performance of the method on the datasets was evaluated and it is shown that MCCP is valid even for severely imbalanced datasets. This indicates that MCCP is a suitable approach for chemogenomics data modelling when an estimation of confidence is desired.

## MONDRIAN CROSS-CONFORMAL PREDICTION

The mathematically formal description and proofs of conformal prediction can be found in the work of Vovk et al.^14^ and detailed descriptions of the conformal prediction framework within the QSAR domain can be found in our previous papers^18-19^. The idea with MCCP is to combine the benefits of a Mondrian conformal predictor and a cross-conformal predictor. Here we will first briefly discuss how conformal prediction estimates the prediction confidence in general. Then the concept of Mondrian conformal prediction will be discussed. We confine our discussion to classification problems only, but a similar approach can be adopted to regression problems.

### Conformal prediction

Intuitively, the problem of conformal prediction is how to estimate the confidence of predicting the class label, *y*, to an new object, *x,* for which given a training set of examples, *z*_1_ = (*x*_1_,*y*_1_), *z*_2_ = (*x*_2_,*y*_2_),…,*z_l_* = (*x_l_*,*y_l_*), the example *z*_i_ = (*x_i_*, *y_i_*) conforms to. This is done by finding how strange (nonconformal) a new example is in comparison to the training set, by calculating the nonconformity measure (NCM) of *z_i_*, assuming each and every possible class label that the object, *x_i_,* can have according to,

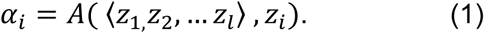

Here the 〈…〉 denotes a bag or a collection of examples and *α_i_* is the NCM of test example *z_i_*. The function *A* is defined by an underlying machine-learning model based on the training set. We remark that the machine learning can be of any type as long as it does not violate the requirement that the training set and any examples that would be predicted are exchangeable. To assess how different a new example *z_i_* is from all old examples, we need to compare *α_i_* to *α_j_* of the previous examples *z_j_* (*j* = 1,…,*l*).

In the inductive learning setting, which would normally be used in QSAR, we can split the training set into a proper training set (*z*_1_*, …,z_m_*) and a calibration set (*z_m+_*_1_, …,*z_l_*), where *m* < *l*. The examples (*z_l+_*_1_,…,*z_l+k_*) are in the prediction set. The p-value for every prediction example *z_i_* can then be calculated as

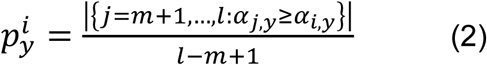

where only calibration set examples are used to calculate p-values in inductive conformal prediction (ICP)^22^ and the proper training set is used to define the NCM. This reduces the computational overhead since only a single model is built from a training set.

For the classification model of conformal prediction, the region prediction Γ^*ϵ*^ for every test object is calculated as

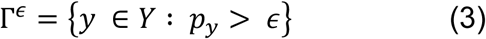

where *Y* is the set of the possible class labels and Γ^*ϵ*^ can be empty or contain one or more classes at a significance level *ϵ.* If the data sets are exchangeable then the predictions will be wrong at a fraction of the number of predictions that will not exceed the significance level. For a binary classifier with the two classes represented by active and inactive, the prediction region could be any of the following sets: {active}, {inactive}, {active, inactive} (both) or {null} (the empty set, i.e. the prediction is that the new example belong neither to the active nor to the inactive class). In this case, a prediction is always considered to be wrong if the set is empty and it is always correct if all possible class labels are predicted.

Several variants of conformal predictions have been proposed^22-24^. One of them is cross-conformal prediction (CCP)^23^ that divides the data into *k* folds the way cross-validation works so that all training data is used as calibration set. Each fold is used once as a calibration set and the remaining training data is used to compute NCMs. For a single prediction, this would lead to *k* p-values being predicted for each possible class label. The final output of those p-values would be to for example report the mean p-value for each possible class label. The motivation for using a CCP is to use all training data for calibration and NCM calculations. However, theoretical guarantees have not been shown in terms of validity.

### Mondrian cross-conformal prediction paradigm

The validity guarantee of the conformal predictor is based on all class labels, not on individual labels. This might be problematic for some applications, in particular if the datasets are imbalanced. For instance, if only 1% of the compounds in a dataset have the label active, most of the active compounds might be assigned as inactive and the conformal prediction would still be considered as valid. Mondrian conformal prediction was developed to address this issue. In the Mondrian framework, the p-value for a hypothesis *y_l_*_+1_ *= y* for the label of test object *z_l_*_+1_ is defined as follows

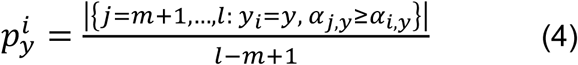

The difference with respect to the definition of p-value in cross-conformal prediction is that the NCM *α_i_* comparisons are restricted to training examples with the same class label. This transforms the global validity guarantee into a label specific guarantee.

Per definition the selection of the calibration set will influence the region prediction in ICP setting. In this investigation a new protocol, called Mondrian cross-conformal prediction (MCCP), is proposed to alleviate the bias caused by randomly selecting the calibration set. The concept is illustrated in Figure 1 and is similar to k-fold cross-validation. The original training sample is randomly partitioned into *k* equal sized subsamples. Of the *k* subsamples, a single subsample is retained as the calibration set for calculating the p-value as in Equation 4, and the remaining *k*− 1 subsamples are used as the proper training set for model building. The process is then repeated *k* times (the *folds),* with each of the *k* subsamples used exactly once as the calibration set. The *k* p-values from the folds can then be averaged to produce a single estimation for the final prediction region of prediction objects.

**Figure 1.**
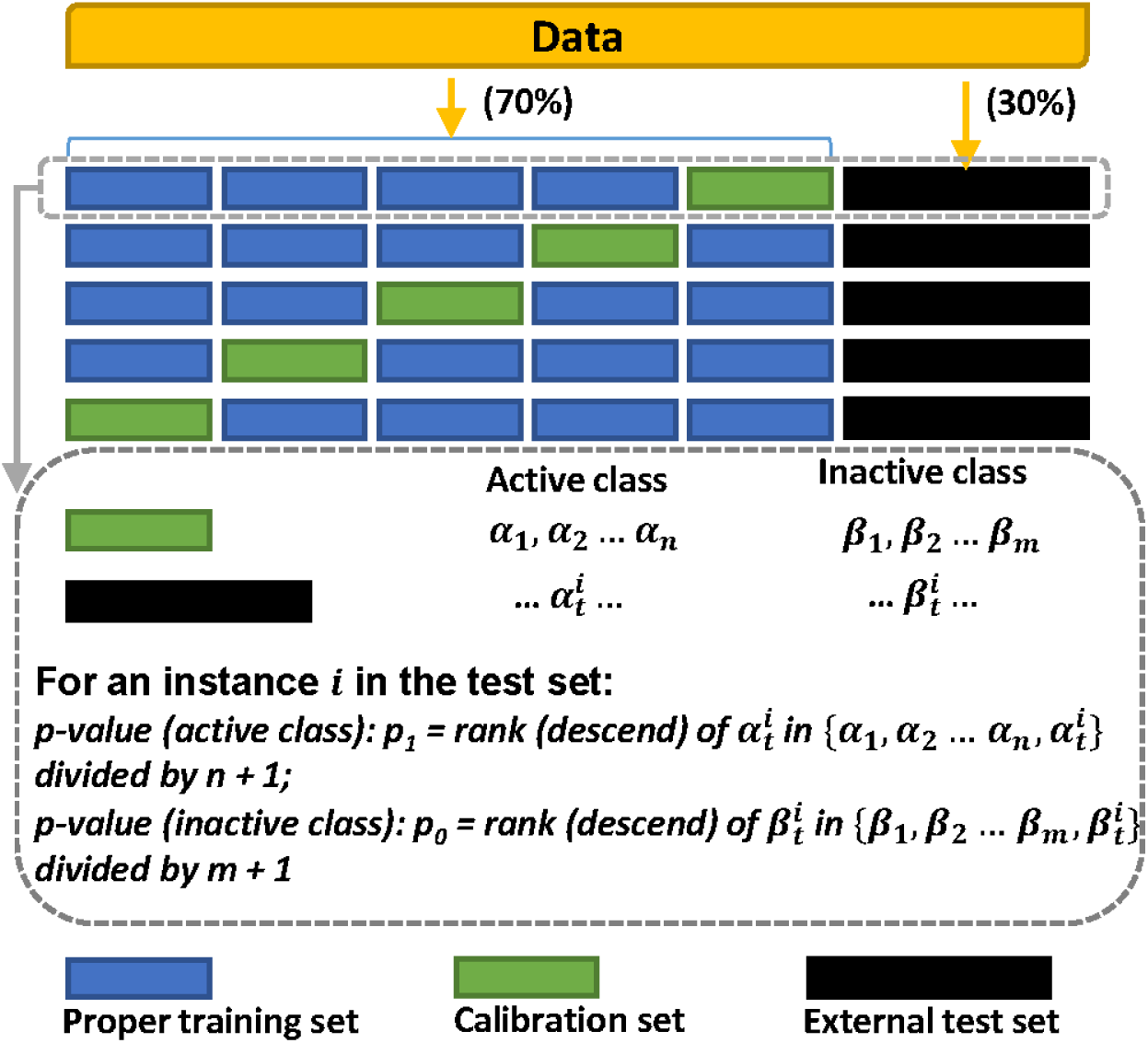
Mondrian cross-conformal prediction framework.

## MATERIAL AND METHODS

### Datasets

18 datasets were extracted from the ExCAPE-DB database^25^, a repository storing public available chemogenomics data. The datasets are binary (active/inactive) and have levels of imbalance varying from 1:10 to 1:1000 (ratio of active/inactive, listed in Table 1). Dataset structures and activity labels are deposited in the GitHub^26^ Signature descriptors^27^ of heights 0-3 were generated for all the compounds.

**Table 1.**
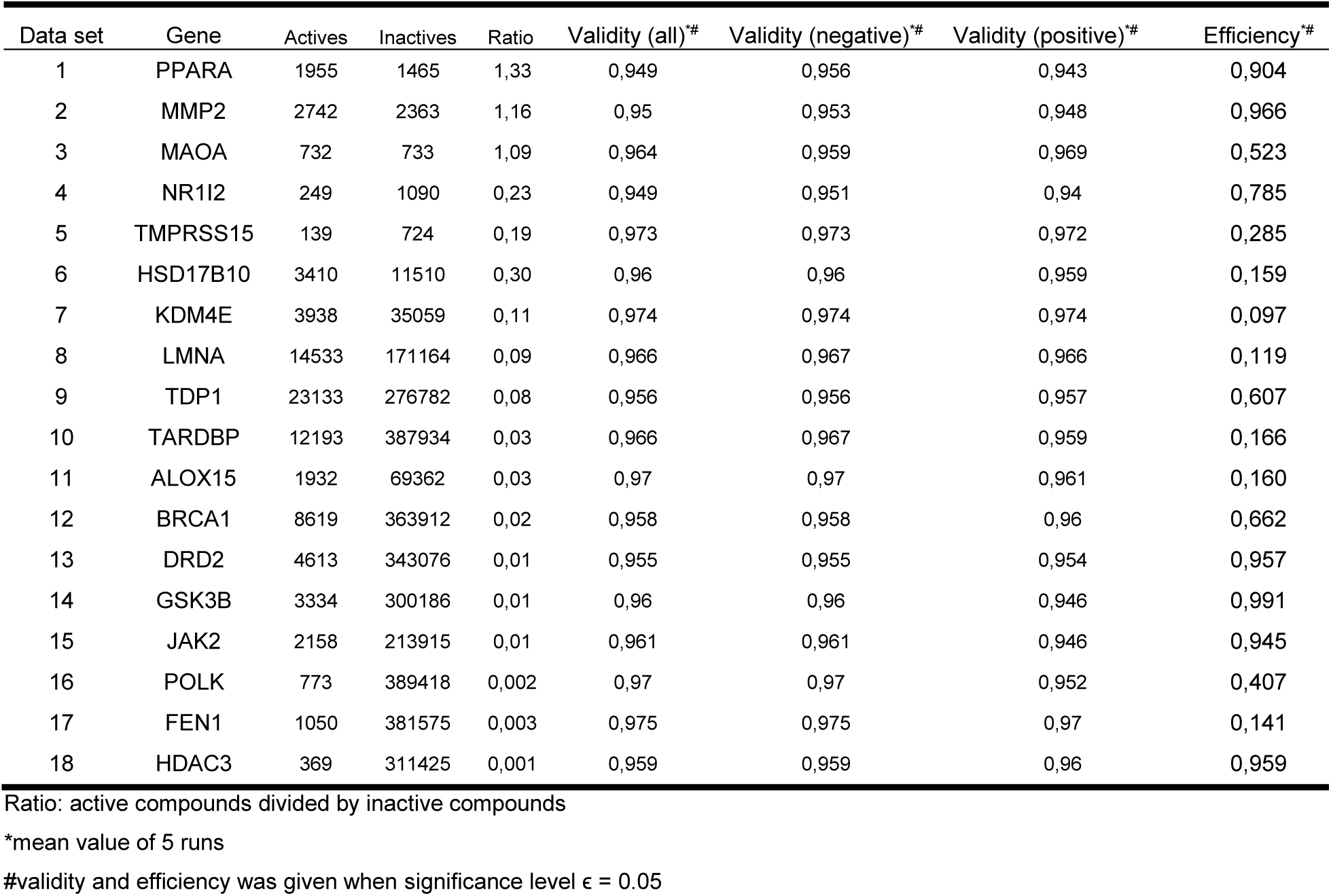
Performance of MCCPs for the 18 datasets.

| Data set | Gene | Actives | Inactives | Ratio | Validity (all)<sup>*#</sup> | Validity (negative)<sup>*#</sup> | Validity (positive)<sup>*#</sup> | Efficiency<sup>*#</sup> |
| --- | --- | --- | --- | --- | --- | --- | --- | --- |
| 1 | PPARA | 1955 | 1465 | 1,33 | 0,949 | 0,956 | 0,943 | 0,904 |
| 2 | MMP2 | 2742 | 2363 | 1,16 | 0,95 | 0,953 | 0,948 | 0,966 |
| 3 | MAOA | 732 | 733 | 1,09 | 0,964 | 0,959 | 0,969 | 0,523 |
| 4 | NR1I2 | 249 | 1090 | 0,23 | 0,949 | 0,951 | 0,94 | 0,785 |
| 5 | TMPRSS15 | 139 | 724 | 0,19 | 0,973 | 0,973 | 0,972 | 0,285 |
| 6 | HSD17B10 | 3410 | 11510 | 0,30 | 0,96 | 0,96 | 0,959 | 0,159 |
| 7 | KDM4E | 3938 | 35059 | 0,11 | 0,974 | 0,974 | 0,974 | 0,097 |
| 8 | LMNA | 14533 | 171164 | 0,09 | 0,966 | 0,967 | 0,966 | 0,119 |
| 9 | TDP1 | 23133 | 276782 | 0,08 | 0,956 | 0,956 | 0,957 | 0,607 |
| 10 | TARDBP | 12193 | 387934 | 0,03 | 0,966 | 0,967 | 0,959 | 0,166 |
| 11 | ALOX15 | 1932 | 69362 | 0,03 | 0,97 | 0,97 | 0,961 | 0,160 |
| 12 | BRCA1 | 8619 | 363912 | 0,02 | 0,958 | 0,958 | 0,96 | 0,662 |
| 13 | DRD2 | 4613 | 343076 | 0,01 | 0,955 | 0,955 | 0,954 | 0,957 |
| 14 | GSK3B | 3334 | 300186 | 0,01 | 0,96 | 0,96 | 0,946 | 0,991 |
| 15 | JAK2 | 2158 | 213915 | 0,01 | 0,961 | 0,961 | 0,946 | 0,945 |
| 16 | POLK | 773 | 389418 | 0,002 | 0,97 | 0,97 | 0,952 | 0,407 |
| 17 | FEN1 | 1050 | 381575 | 0,003 | 0,975 | 0,975 | 0,97 | 0,141 |
| 18 | HDAC3 | 369 | 311425 | 0,001 | 0,959 | 0,959 | 0,96 | 0,959 |

Ratio: active compounds divided by inactive compounds
\*mean value of 5 runs

### Application of inductive conformal prediction

Both MCCP and CCP were performed on all 18 datasets to compare their performance. The MCCP workflow is displayed in Figure 1. First, the dataset was randomly divided into two parts: training (70%) and external test (30%) set. As described earlier, the training set was randomly partitioned into 5 folds for estimating prediction regions of test set compounds using MCCP. The support vector machine (SVM) module of the Scikit-Learn^28^ was used to build SVM models. We defined the NCM by using the SVM decision function as follows:

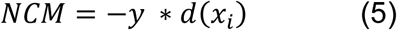

where *y* is the non-zero class label (1, −1) and *d*(*x_i_*) is the decision value obtain from the SVM decision function for compound *x_i_*.

For comparison, the Tanimoto distance between a specific compound and the proper training set was also used as a NCM. The distance was calculated by averaging the Tanimoto similarity values between the five most similar compounds in the proper training set and the specific compound. The pairwise Tanimoto similarity was calculated in Scikit-Learn^28^ using the 2048-length bit string Extended-Connectivity Fingerprints (ECFPs).

The jCompoundMapper^29^ was used to generate ECFP fingerprints by setting search depth to 6.

### Evaluation metrics

For a compound, given that the p-value for active and inactive class is *p*_1_ and *p_0_* respectively, an output label from conformal prediction under significance level ϵ can be defined as following:

Active: *p*_1_ > *ϵ* and *p*_0_ ≤ *ϵ*

Inactive: *p*_0_ > *ϵ* and *p*_1_ ≤ *ϵ*

Uncertain (Both): *p*_1_ > *ϵ* and *p*_0_ > *ϵ*

Empty (None): *p*_1_ *< ϵ* and *p*_0_ ≤ *ϵ*

Two measurements, validity and efficiency, are used to measure the performance of conformal prediction. The conformal prediction is said to be valid if the frequency of errors (i.e., the fraction of true values outside the prediction region) is less than ϵ at a chosen confidence level 1 − *ϵ.* The validity can be calculated for all class objects as well as for objects of one specific class. Efficiency is defined as the observed singleton prediction set rate at a given significance value *ϵ*^30^. Here, singleton means either predicted as active or as inactive.

Cohen's kappa^31^ is a classification metrics designed to measure the agreement between the observed and the predicted labels of test set according to equation below

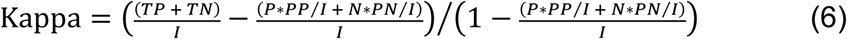

where TP denotes the number of true positive, TN the number of true negative, P the number of positive instances, N the number of negative instances, PP the number of the predicted positives, PN the number of the predicted negatives, and I the number of total instances.

## RESULTS AND DISCUSSION

### Performances of Mondrian cross-conformal prediction

The validity curves in Figure 2A shows that MCCPs in general are valid for both balanced and imbalanced datasets for significance values less than 0.2 and also confirmed in Table 1 at significance level 0.05 are all lower than 0.05. The validity is also demonstrated for both active and inactive classes. Notably, MCCPs are also valid for the minority class of very imbalanced datasets, e g. data set **18**. The observed singleton prediction set rates (efficiency) are good for the balanced (data sets **1** and **2**) and some highly imbalanced dataset (data sets **13-15** and **18**) but lower for data sets **6-8**, **10-11** and **17** when the significance value õ is less than 0.2.

**Figure 2.**
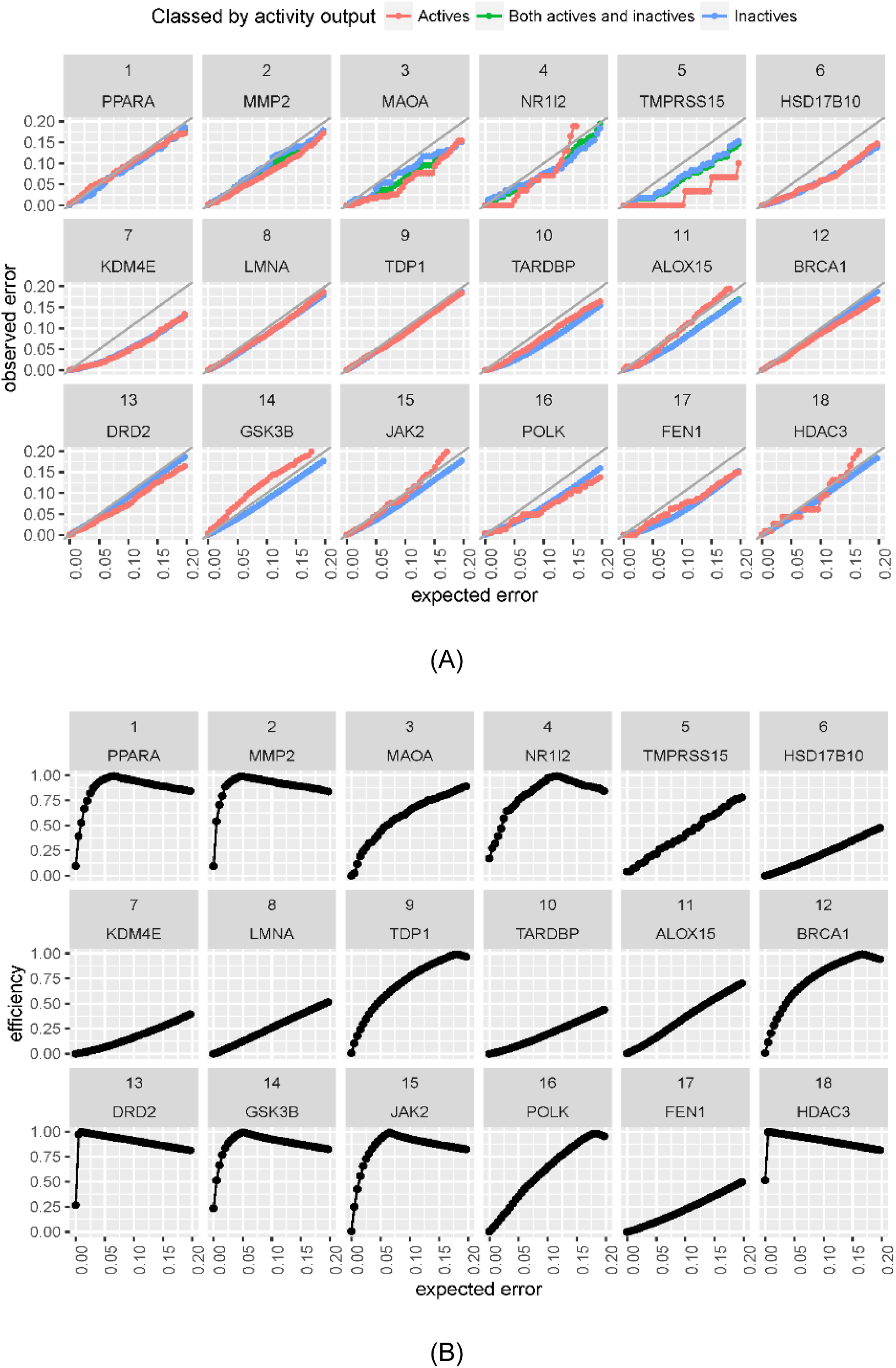
Performances of MCCP. (A) Validities and (B) efficiencies of the18 data sets. Active (red), inactive (blue) and both classes (green) are displayed separately. The colour of curves corresponds to different type of examples and the light grey refers to the diagonal line to demonstrate the validity. Validities in different class may have overlaps in some data sets that make some colours invisible

### Performances of cross-conformal prediction

As a comparison to MCCP, the CCP method was applied on the same datasets (Figure 3A-B and Table 2). It can be seen that the global validity of the CCP models is achieved on both balanced and imbalanced datasets. Investigating the local validity for each class label, the CCP models are still valid for the balanced dataset (e g. Data sets **1-3**). However, the CCP models do not seem to be valid for the minority class in the imbalanced datasets (e.g. data sets **4-18**) whose active-to-inactive-ratios are less than 1:5. It is likely due to that CCP tends to predict the active data points (minority class) as inactive (majority class) in those data sets. But CCP models generally have higher efficiency compared to that of MCCPs. These results demonstrate, as previously discussed, that CCP models cannot guarantee the label-conditional validity for all labels on imbalanced datasets, while MCCPs can obtain both the global and label-conditional validity even for highly imbalanced datasets.

**Figure 3.**
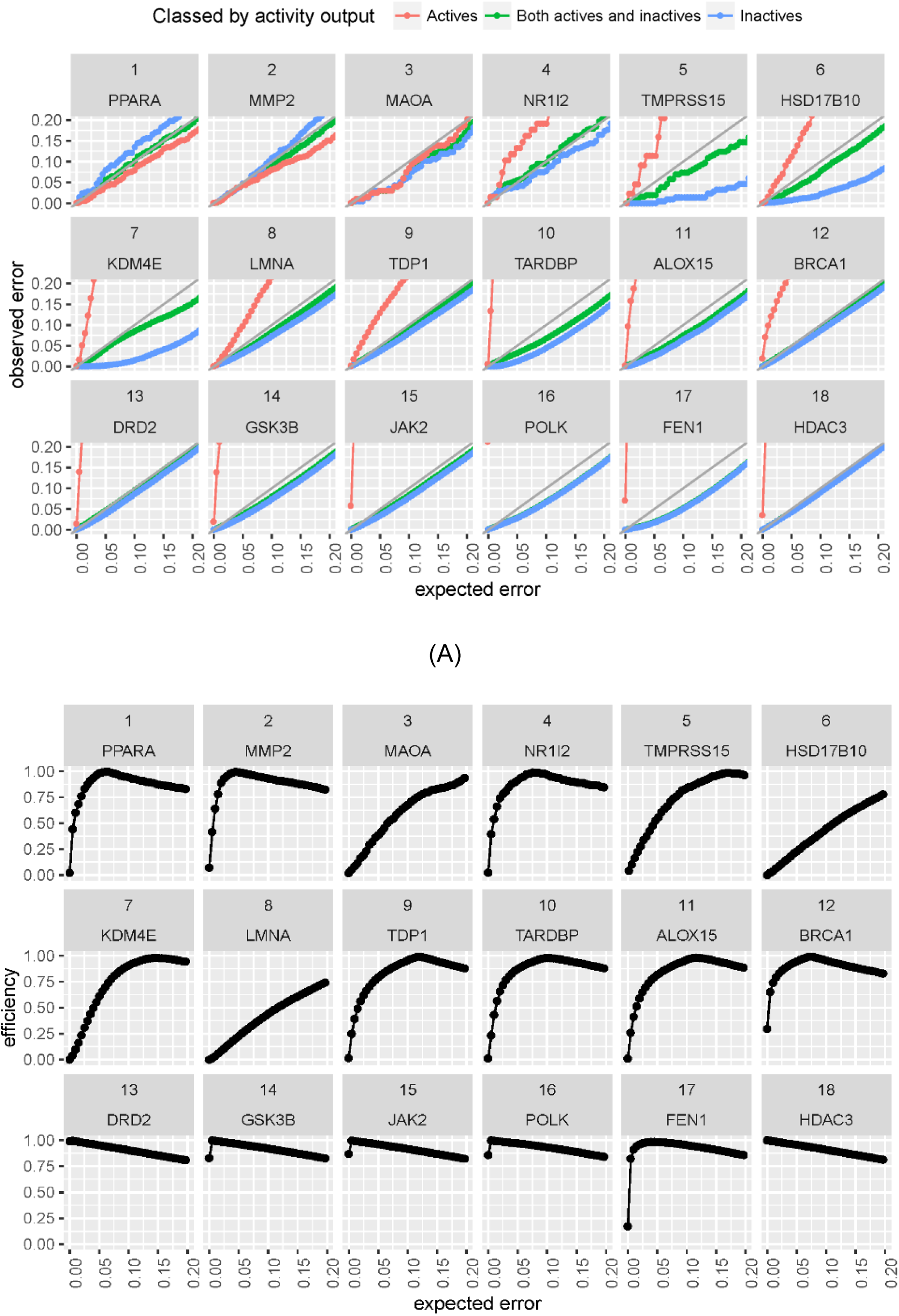
Performances of CCP. (A) Validities and (B) efficiencies of 18 data sets. Active (red), inactive (blue) and both classes (green) are displayed separately. Validities in different class may have overlaps in some data sets that make some colours invisible.

**Table 2.** Performance of CCPs for the 18 datasets.

| Data set | Gene | Validity (all)<sup>*#</sup> | Validity (negative)<sup>*#</sup> | Validity (positive)<sup>*#</sup> | Efficiency<sup>*#</sup> |
| --- | --- | --- | --- | --- | --- |
| 1 | PPARA | 0,949 | 0,932 | 0,962 | 0,979 |
| 2 | MMP2 | 0,957 | 0,951 | 0,963 | 0,994 |
| 3 | MAOA | 0,96 | 0,965 | 0,954 | 0,461 |
| 4 | NR1I2 | 0,936 | 0,951 | 0,867 | 0,911 |
| 5 | TMPRSS15 | 0,963 | 0,993 | 0,804 | 0,654 |
| 6 | HSD17B10 | 0,969 | 0,994 | 0,884 | 0,232 |
| 7 | KDM4E | 0,955 | 0,998 | 0,581 | 0,624 |
| 8 | LMNA | 0,962 | 0,968 | 0,896 | 0,232 |
| 9 | TDP1 | 0,956 | 0,963 | 0,868 | 0,797 |
| 10 | TARDBP | 0,967 | 0,988 | 0,313 | 0,879 |
| 11 | ALOX15 | 0,967 | 0,976 | 0,615 | 0,830 |
| 12 | BRCA1 | 0,955 | 0,96 | 0,769 | 0,947 |
| 13 | DRD2 | 0,954 | 0,962 | 0,362 | 0,954 |
| 14 | GSK3B | 0,962 | 0,967 | 0,456 | 0,963 |
| 15 | JAK2 | 0,963 | 0,967 | 0,483 | 0,963 |
| 16 | POLK | 0,97 | 0,971 | 0,25 | 0,971 |
| 17 | FEN1 | 0,98 | 0,982 | 0,31 | 0,985 |
| 18 | HDAC3 | 0,957 | 0,958 | 0,412 | 0,957 |

\*mean value of 5 runs
#validity and efficiency was given when significance level $\epsilon = 0.05$

### Comparison of accuracy between MCCP and ordinary SVM prediction

Although the main goal of the MCCP method is to provide confidence estimation for prediction, it is still interesting to investigate if MCCP can provide better prediction at certain significance levels than the ordinary SVM prediction. Therefore, SVM calculations is done using the RBF kernel on the same data set as MCCP. The proper training and calibration set is merged together as the SVM training set. We compare the prediction performances of MCCP and ordinary SVM in terms of Kappa. To make a fair comparison, only instances which have a singleton class prediction in MCCP were used for computing the Kappa value for the SVM model (Table 3 and Figure 4). At significance level ϵ of 0.05 (i.e. corresponding to confidence level of 95%), MCCPs and SVMs have almost the same accuracy in most cases based on Kappa. This is logical since that the same underlying classifier is used for both MCCP used SVM. It was also noticed that SVM has better performance on data sets **14**-**16** at 0.05 significance level, which might be due to that the actual training set is larger. Nevertheless, MCCP can in general achieve the same level of accuracy as ordinary SVM while provide additional confidence estimations. Moreover, MCCP outperformed SVM in most cases except for data sets **13-16** and **18** if the kappa values of SVM were computed based on all compounds in the test set (Figure 4).

**Table 3.** Performance of MCCP versus SVM.

| Data set | Gene | Kappa<sup>#</sup> (M CCP) | Kappa<sup>#</sup> (SVM) | Kappa<sup>†</sup> (SVM) |
| --- | --- | --- | --- | --- |
| 1 | PPARA | 0,887 | 0,888 | 0,880 |
| 2 | MMP2 | 0,899 | 0,899 | 0,912 |
| 3 | MAOA | 0,861 | 0,861 | 0,568 |
| 4 | NR1I2 | 0,812 | 0,813 | 0,776 |
| 5 | TMPRSS15 | 0,752 | 0,752 | 0,522 |
| 6 | HSD17B10 | 0,376 | 0,376 | 0,223 |
| 7 | KDM4E | 0,383 | 0,383 | 0,175 |
| 8 | LMNA | 0,227 | 0,227 | 0,093 |
| 9 | TDP1 | 0,624 | 0,624 | 0,451 |
| 10 | TARDBP | 0,145 | 0,145 | 0,081 |
| 11 | ALOX15 | 0,286 | 0,328 | 0,160 |
| 12 | BRCA1 | 0,366 | 0,381 | 0,287 |
| 13 | DRD2 | 0,933 | 0,933 | 0,938 |
| 14 | GSK3B | 0,322 | 0,761 | 0,784 |
| 15 | JAK2 | 0,298 | 0,725 | 0,831 |
| 16 | POLK | 0,064 | 0,314 | 0,268 |
| 17 | FEN1 | 0,050 | 0,126 | 0,049 |
| 18 | HDAC3 | 0,918 | 0,917 | 0,845 |

\*mean value of 5 runs
#value was given when significance level $\epsilon = 0.05$ , only prediction of singleton class in M CCP were considered
†value was computed based on the whole test set

**Figure 4.**
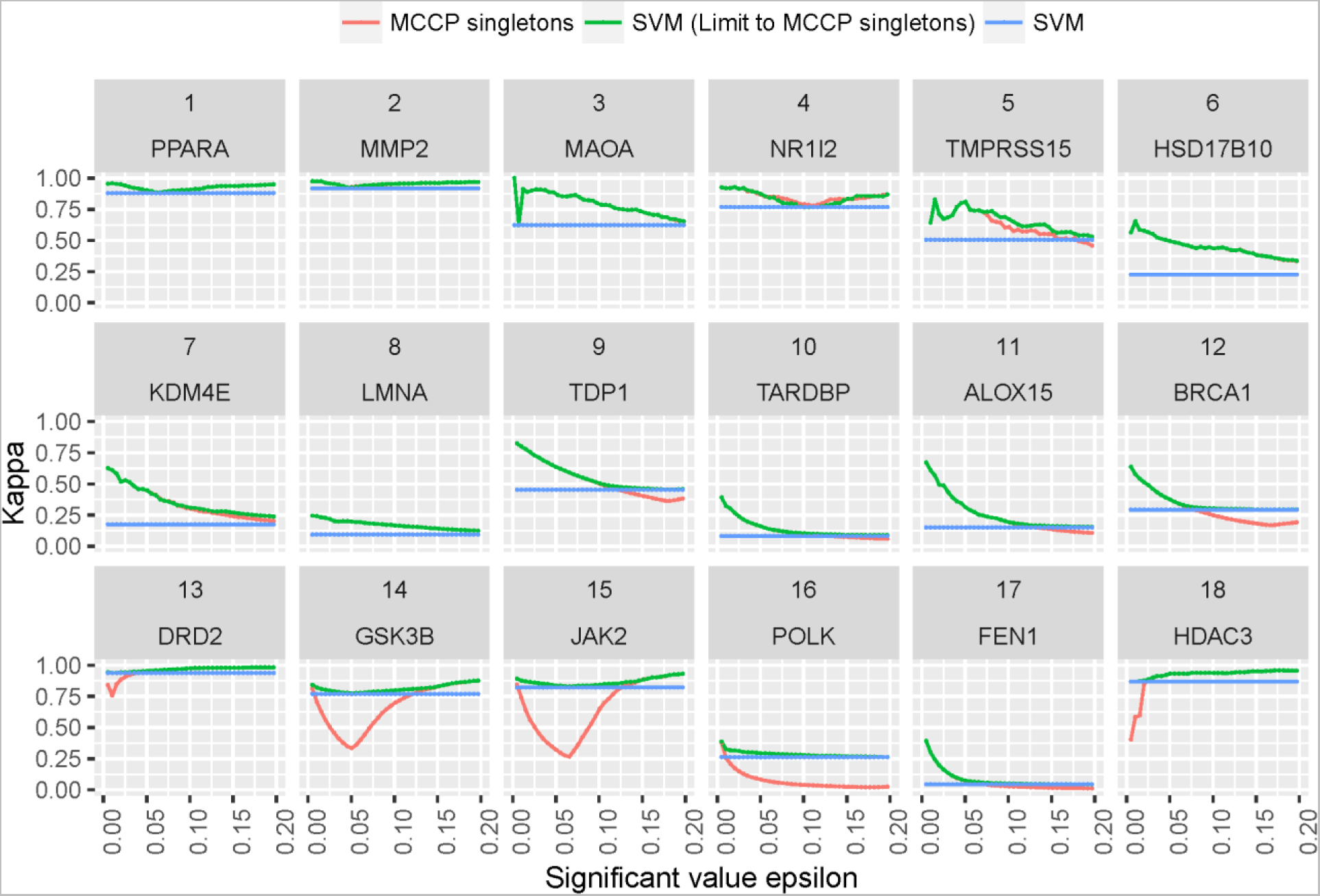
Comparison of performance between MCCP and ordinary SVM. Kappa values are computed based on singletons of MCCP (red) and its corresponding sets in SVM (green), and all instances in SVM (blue), respectively. Kappa values in different group may have overlaps in some data sets that make some colours invisible.

### Influence of different nonconformity measure

The NCM function is used in conformal prediction framework to characterise the nonconformity between a test example and the training examples. Per definition, most AD measures can be easily adopted as NCM and integrated into the conformal prediction paradigm. Calculating similarity between test compound and its k nearest neighbours (k-NN) among training compounds is usually used as an intuitive way of measuring AD^32^. Here we compare the region prediction performance of k-NN similarity based NCM and the default NCM using the SVM embedded decision function. For the k-NN based NCM calculation, the top five most similar compounds in the training set were obtained for each query (test) compound and the NCM is the average similarity distance to the five nearest neighbours.

The validity and efficiency plots of k-NN based MCCP models are shown in Figure 5A-B. MCCPs based on k-NN similarity are also valid for most datasets (except data sets **4** and **5**) but their efficiency are lower than 0.4, which means for most of compounds (more than 60% of compounds in the test set) that the prediction region is either empty or uncertain. In contrast, a higher efficiency can be obtained when the NCM is based on the SVM decision value (Figures 2B and 3B) compared to that based on k-NN similarity. This demonstrate that k-NN similarity is a valid but not an efficient NCM. This is not surprising since the k-NN similarity is a model independent metrics generated in a non-supervised manner, while the SVM decision value is a model dependent metrics which has been optimized on the training set.

**Figure 5.**
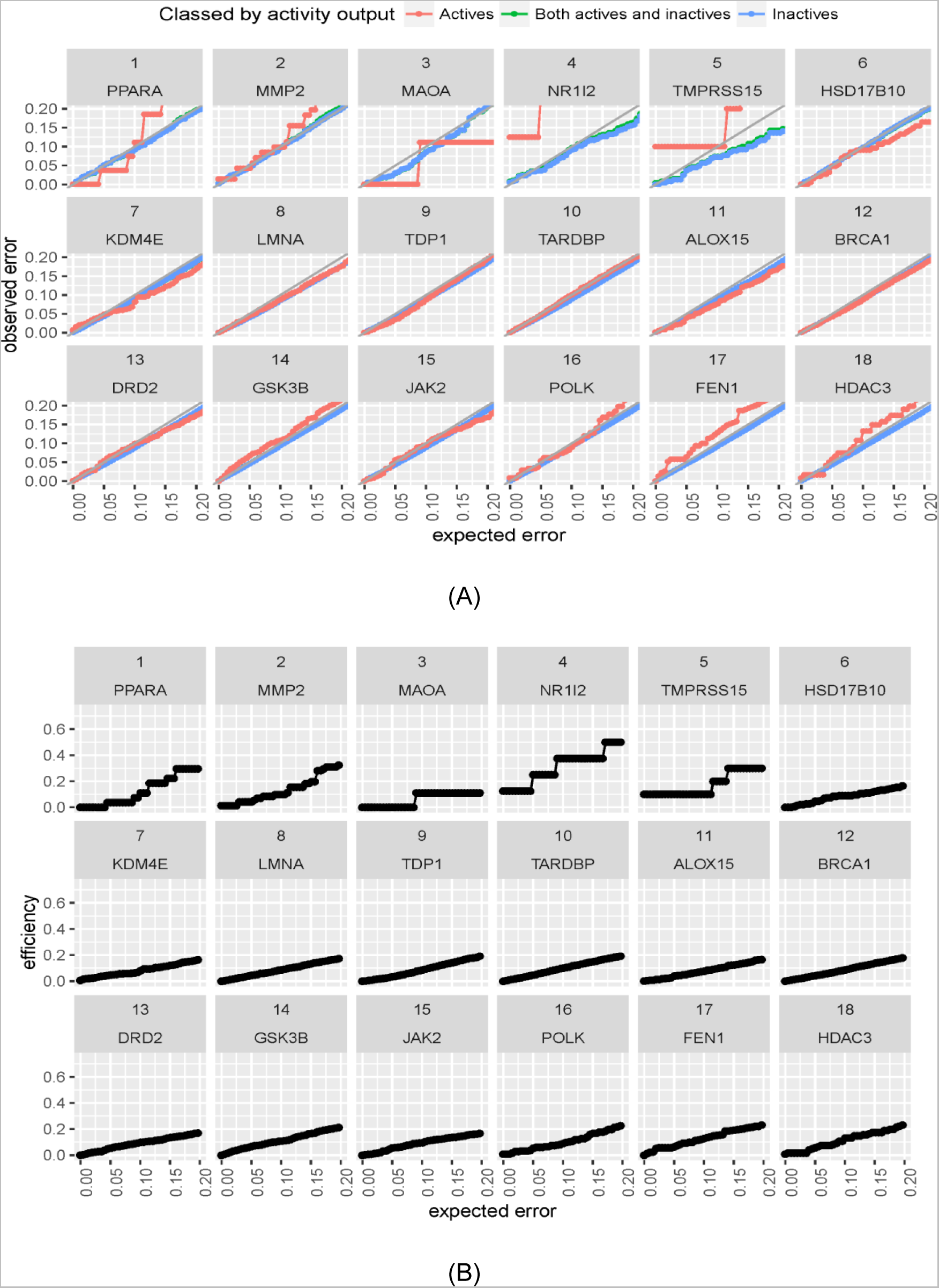
Performance of MCCP based on k-NN. (A) Validities and (B) efficiencies of 18 data sets. Active (red), inactive (blue) and both classes (green) are displayed separately. Validities in different class may have overlaps in some data sets that make some colours invisible.

## CONCLUSIONS

In this study, the conformal prediction protocol MCCP was for the first time used for an application in cheminformatics. We investigated the validity and efficiency of MCCP on 18 large scale cheminformatics datasets with various levels of imbalance and compared the method with conventional CCP. Our results show that the MCCP confidence estimation is valid globally, i.e. overall for both classes, as well as locally for each class. While CCP models is not guaranteed to be valid for the minority class for imbalanced data sets. The prediction accuracy of MCCP model is similar to the original SVM model at significance level 0.05. The k-NN similarity based NCM is also evaluated in MCCP and compared with the SVM decision value based NCM. Although the k-NN similarity based confidence estimation is valid for most of the data sets, its efficiency is significantly worse than the SVM decision value based NCM. This result highlights the importance of choosing a suitable NCM with respect to the efficiency of conformal prediction.

## ACKNOWLEDGMENTS

This research has received funding from the ExCAPE project within European Union’s Horizon 2020 framework under Grant Agreement no. 671555. We thank Prof. Alex Gammerman and Dr. Paolo Toccaceli (Royal Holloway, University of London) for helpful discussions.

The research at Swetox (UN) was supported by Stockholm County Council, Knut & Alice Wallenberg Foundation, and Swedish Research Council FORMAS.

